# A High Resolution Map of Meiotic Recombination in *Cryptococcus* Demonstrates Decreased Recombination in Unisexual Reproduction

**DOI:** 10.1101/234617

**Authors:** Cullen Roth, Sheng Sun, R. Blake Billmyre, Joseph Heitman, Paul M. Magwene

## Abstract

Multiple species within the basidiomycete genus, *Cryptococcus*, cause cryptococcal disease. These species are estimated to affect nearly a quarter of a million people leading to approximately 180,000 mortalities, annually. Sexual reproduction, which can occur between haploid yeasts of the same or opposite mating type, is a potentially important contributor to pathogenesis as recombination can generate novel genotypes and transgressive phenotypes. However, our quantitative understanding of recombination in this clinically important yeast is limited. Here we describe genome-wide estimates of recombination rates in *Cryptococcus deneoformans* and compare recombination between progeny from α-α unisexual and **a**-α bisexual crosses. We find that offspring from bisexual crosses have modestly higher average rates of recombination than those derived from unisexual crosses. Recombination hot and cold spots across the *C. deneoformans* genome are also identified and are associated with increased GC content. Finally, we observed regions genome-wide with allele frequencies deviating from the expected parental ratio. These findings and observations advance our quantitative understanding of the genetic events that occur during sexual reproduction in *C. deneoformans*, and the impact that different forms of sexual reproduction are likely to have on genetic diversity in this important fungal pathogen.

---

Annually, cryptococcal disease is estimated to affect more than 200,000 people worldwide, accounting for approximately 15% of AIDS-related mortalities (Rajasingham *et al.* 2017). While *Cryptococcus* species are preferentially haploid (Hull *et al.* 2002) and propagate primarily asexually, sexual reproduction and recombination have been demonstrated in both the laboratory and environment (Kwon-Chung 1975,1976; Litvintseva *et al.* 2003; Lin *et al.* 2007; Hull *et al.* 2002). The sexual cycle in *Cryptococcus* has clinical relevance as sexual reproduction produces spores, which can serve as infectious propagules, that are readily aerosolized and inhaled by hosts (Giles *et al.* 2009; Velagapudi *et al.* 2009; Coelho *et al.* 2014). Furthermore, recombination during sex produces new genotypes, some of which may display novel phenotypes linked to virulence, such as the ability of offspring to grow at higher temperatures than that of their parental strains (Sun *et al.* 2014). Thus, quantitatively characterizing recombination in *Cryptococcus* is a key step to developing a better understanding of the genetics of virulence in this clade.

*Cryptococcus deneoformans* (formerly *C. neoformans* var. *neoformans* (serotype D), see Hagen *et al.* (2015) and Kwon-Chung *et al.* (2017) for recent discussions of nomenclature) possesses a bipolar mating system with the mating type locus (*MAT*) on the fourth chromosome. The *MAT* locus, which is greater than 100 kb in size and contains more than 20 genes, is represented in two mating type alleles, α and a (Heitman *et al.* 1999; Lengeler *et al.* 2002; Loftus *et al.* 2005; Sun and Heitman 2016). In the laboratory setting, sexual reproduction has been observed between haploid *MAT*α and *MAT***a** strains (Kwon-Chung 1976; Hull *et al.* 2002; Xue *et al.* 2007; Nielsen *et al.* 2007; Sun *et al.* 2014; Gyawali *et al.* 2017). Diploid strains and signatures of re-combination have been documented in environmental isolates, indicating that sexual reproduction also occurs in nature (Litvintseva *et al.* 2003; Campbell *et al.* 2005; Lin *et al.* 2007; Bui *et al.* 2008; Lin *et al.* 2009). However, an analysis of environmental and clinical isolates of *Cryptococcus* species revealed a bias in the distribution of the mating type alleles, with the majority of *C. deneoformans* isolates analyzed possessing the *MAT*α allele (Kwon-Chung and Bennett 1978). This observation called into question the frequency and importance of bisexual reproduction, and thus recombination, in the wild. Lin *et al.* (2009) provided an answer to this conundrum with the discovery that *C. deneoformans* is also capable of undergoing same sex or unisexual matings between *MAT*α strains (Lin *et al.* 2005, 2007, 2009).

Meiosis is an integral component of sexual reproduction (Page and Hawley 2003) that occurs in both unisexual and bisexual reproduction (Feretzaki and Heitman 2013). Within a basidium, meiosis produces nuclei that will undergo several rounds of mitosis to generate subsequent nuclei that are packaged into spores (Kwon-Chung 1980). These basidiospores then bud from the basidium in four long chains (Kwon-Chung 1980; Idnurm 2010). Dissection of basidiospore chains and analysis of their genotypes shows segregation of alleles consistent with one round of meiosis and demonstrates that post-meiotic nuclei undergo mitosis and randomly assort into different spore chains (Kwon-Chung 1980; Idnurm 2010).

Various studies have examined recombination rates in *Cryptococcus* species, as well other phenomena that occur during meiosis, such as crossover hot spots, gene conversions, and allele segregation distortion (Forche *et al.* 2000; Marra *et al.* 2004; Hsueh *et al.* 2006; Sun and Xu 2007; Sun *et al.* 2014; Sun and Heitman 2016). Genome-wide, our quantitative understanding of recombination is limited to a few studies of *C. deneoformans* crosses (Forche *et al.* 2000; Marra *et al.* 2004) and hybrid crosses, between *C. deneoformans* and *C. neoformans* strains (Sun and Xu 2007). Current estimates of recombination rates for *C. deneoformans* are based on linkage maps constructed via a modest number of genetic markers, with estimates varying between 13.2 kb/cM (Marra *et al.* 2004) and 7.13 kb/cM (Sun *et al.* 2014), with no observed difference in recombination rates between progeny derived from unisexual versus bisexual reproduction (Sun *et al.* 2014).

In the present study we utilize whole genome sequencing data to quantitatively analyze differences in genome-wide recombination rates between progeny from unisexual and bisexual reproduction, to identify recombination hot and cold spots, and to identify chromosomal regions that exhibit biased or distorted allele frequencies. We find genome-wide differences in the average rates of recombination between progeny from α-α unisexual and **a**-α bisexual crosses, with higher rates of crossovers in samples from **a**-α bisexual crosses. Recombination hot and cold spots are identified, with hot spots associated with higher than average GC content, and cold spots clustering near centromeres. Centromeric cold spots are often flanked by areas of increased crossover activity. Finally, we show that regions with allele frequencies deviating from the expected 2:2 parental allele ratio are not unique to chromosome four and are seen genome-wide. The high resolution characterization of patterns and rates of recombination that this study provides helps to advance our understanding of the processes that generate genetic diversity in this fungus, and will serve as a foundation for future investigations of the population and quantitative genetics of *C. deneoformans* and related *Cryptococcus* species.

## Materials and Methods

### Strains, laboratory crosses and isolation

As previously described (Sun *et al.* 2012, 2014), parental strains XL280**a**, XL280αSS, and 431α, were used in **a**-α bisexual and α-α unisexual crosses. XL280αSS is an XL280α strain with a ectopically inserted *NAT* resistance marker in the *URA5* gene and congenic to XL280**a** with the exceptions of the *URA5* and *MAT* loci. Inverse PCR conducted on the XL280αSS strain confirmed the insertion site of the *NAT* resistance marker within the *URA5* gene. For **a**-α bisexual crosses between strains XL280**a** and 431α, chains of basidiospores from individual basidia were transferred onto fresh YPD medium, and individual basidiospores were separated using a fiber optic needle (Sun *et al.* 2014). From α-α unisexual crosses between XL280αSS and 431α, recombinant progeny were generated using selectable markers to isolate *NAT^R^ URA*^+^ progeny (Sun *et al.* 2014).

### Sequencing, aligning, variant calling and filtering

From the α-α unisexual and the **a**-α bisexual crosses, 105 segregants were isolated for whole genome sequencing (Sun *et al.* 2014). Sequencing was performed on the Illumina Hiseq 2500 platform at the University of North Carolina Chapel Hill Next Generation Sequencing Facility. A paired end library with approximately 300 base inserts was constructed for each sample, and libraries were multiplexed and run 24 samples per lane using 100 bp paired-end reads. Raw reads were aligned to an XL280 *C. deneoformans* reference genome (McKenna *et al.* 2010; Sun *et al.* 2014) using BWA (v0.7.10-r789, Li and Durbin 2009). Variant calling was carried out using The Genome Analysis Toolkit (v3.1-1, McKenna *et al.* 2010) and SAMtools (v1.2, Li 2011) resulting in 143,812 variable sites across the 105 segregants. These sites were scored as 0 or 1 if inherited form the XL280αSS (or XL280**a**) or the 431α parental strains, respectively. Variable sites were filtered on read depth and quality. Across segregants, variable sites were required to have greater than 15× coverage, a quality score, normalized by read depth, of greater than or equal to 20, and a minor allele frequency per site of at least 1%. Only sites with 100% call rate were used in analysis. Variant calls were further filtered to include only sites exhibiting biallelic, single nucleotide polymorphisms, yielding a final total of 86,767 sites.

### Segregant filtering

Read count data for each SNP site was used to screen each of the 108 segregants for gross aneuploidy of chromosomes. In total six segregants were removed due to partial or complete aneuploidy. Aneuploidy of chromosome one was detected in three segregants, a duplication of the right arm of chromosome seven in one segregant, and aneuploidy of chromosome ten in two segregants. For all samples pairwise genetic correlations were calculated to identify pairs of segregants that were genetically identical. These duplicates were removed from analysis to avoid biasing estimates of recombination by sampling a genotype more than once. In total, four pairs of segregants were identified as genetically identical. From each of the four pairs of segregants, one was removed from analysis. One segregant from the **a**-α bisexual cross, SSB593, showed no recombination across the genome except on chromosome four. All of the other chromosomes in the segregant were inherited from the XL280**a** parental strain. This segregant was removed from further analysis. After passing these filtering criteria, 94 segregants, 55 from α-α unisexual crosses and 39 from **a**-α bisexual crosses, were retained for analysis.

### Haplotype construction and filtering

For each sample, SNP data was used to estimate regions with consecutive SNPs inherited from one parent (i.e haplotypes) between XL280**a**, XL280αSS, and 431α. A “minimum” run approach based on inter-marker intervals was used to determine the size of haplotypes (Mancera *et al.* 2008). Briefly, for a set of SNPs within a haplotype with positions *v*_0_, *v*_1_, …*v_n_* along a chromosome, the size of the haplotype in nucleotide bases or length of the intra-marker interval is calculated as *h* = *v_n_* − *v*_0_ + 1. The inter-marker interval is defined as the distance between two SNPs with opposing genotypes (Mancera *et al.* 2008). Let *v*, *w* be the positions of two adjacent SNPs along a chromosome with opposing genotypes, then the distance in nucleotide bases between the two SNPs is calculated as *d* = *w* − *v* – 1. For each sample, SNP data was used to construct haplotype blocks, where runs of contiguous SNPs with shared genotypes are grouped. For the results shown here, haplotypes were retained if the size of the haplotype or intra-marker interval was greater than or equal to 6 kb.

### Crossover frequency estimation

#### Poisson regression

Haplotype data for each segregant was used to calculate the number of crossovers. For any given segregant with *n* haplotypes there are *n* − 1 crossovers. Genome-wide recombination rates were estimated using Poisson regression, modeling the number of crossovers as a function of chromosome length with the mode of sexual reproduction as a co-variate using the “glm” function implemented in R (version 3.4.1). Our analysis indicated no support for an interaction term between chromosome length and mode of sexual reproduction; we therefore fit a simple additive model of the form log(E(# of crossovers | x)) = *β*_0_ + *β*_1_ **I**_*c*_ + *β*_2_x, where x is chromosome length and **I**_*c*_ is an indicator variable for the cross type (0 = α-α unisexual, 1 = **a**-α bisexual crosses).

The model was estimated as: log(E(# of crossovers | x)) = −0.015 + 0.274**I**_*c*_ + 0.570x. The model fit failed to reject the null hypothesis of a zero intercept term (*B*_0_) but there was strong support to reject the null hypothesis of zero valued *β*_1_ and *β*_2_ coefficients (p-values < 10^−10^).

#### Analysis of crossovers per chromosome

For each chromosome, the number of crossovers was compared between segregants from the α-α unisexual and **a**-α bisexual crosses. A two sided, Mann-Whitney U-test with an *α* = 0.05 was utilized to test for significant differences in the average number of crossovers (per chromosome) along with the Holm-Sidak step down method to correct for multiple testing (Holm 1979).

### Crossover hot and cold Spot discovery and analysis

#### Statistical association testing

For each chromosome, contiguous bins of 41.5 kb were constructed, tiling each chromosome from the edges of the centromeres out to the ends of the chromosome (centromeric regions were excluded from hot/cold spot analysis). After investigating the total detected number of hot and cold crossover spots as a function of bin size (from 0.5 to 100 kb), a bin size of 41.5 kb was chosen because it minimized the difference between the detected number of crossover hot spots and crossover cold spots. The outermost 5′ and 3′ bins of each chromosome were constructed to have at least half of their width overlap the last two annotated SNP on the respected end of that chromosome. Within each bin, the number of inter-marker intervals in which a crossover was detected were counted. For each inter-marker interval, crossovers shared by meiotic siblings were only counted once. For every bin, a Poisson model, with parameters established from genome-wide analysis of crossover frequencies of meiotic progeny from the **a**-α bisexual crosses, was utilized to compare the number of crossovers observed versus the number expected given the bin size. A two-tailed test was used to search for statistically cold and hot crossover spots. A false discovery rate approach (Benjamini and Yekutieli 2001) was used to define genome-wide, significantly hot or cold crossover spots, using at a false-discovery rate cutoff of 0.05. An “artificial” hot spot on chromosome seven, resulting from the use of selectable markers to isolate recombinant progeny from the α-α unisexual crosses (Sun *et al.* 2012, 2014), was removed from the analysis.

#### Analysis of GC content

For each inter-marker interval, nucleotide sequences were obtained from the XL280 reference genome. The GC content for all inter-marker intervals was calculated and classified as hot, cold, or other according to whether the interval fell within a hot or cold region as defined above. In total there were 7,558 hot inter-marker interval sequences, 7,369 cold spot inter-marker interval sequences, and 68,051 intervals defined as other. The GC content for intermarker intervals within hot and cold spots was compared using a two-sided, Mann-Whitney U test (*α* = 0.05). For the three groups of inter-marker interval sequences, 95% confidence intervals were calculated via permutation (sampling with replacement), taking the difference between the observed mean GC content and the sampled mean, 1,000 times. From these deviations, the 2.5% and 97.5% percentiles of the permuted distribution were used as critical values.

#### Identification of motifs associated with crossover hot spot sequences

To search for sequence motifs associated with hot spots, 100 random sequences from hot spot inter-marker intervals in which there was a crossover where chosen such that the lengths of sequences ranged between 100 and 10,000 bases and the sum of the sequences was less than 60 kb (constraints related to the online MEME tool). A complementary control set of 100 randomly chosen sequences were selected from other genomic regions using the same parameters. The hot and control sets of sequences where analyzed using MEME, version 4.12.0 (Bailey *et al.* 1994). Analysis in MEME was conducted using discriminative mode, with zero or one occurrence of a contributing motif site per sequence, searching for four motifs between six and 50 bases wide.

### Analysis of allele distortion and bias

#### Segregants used in haplotype analysis

From the **a**-α bisexual cross, 22 of the 39 segregants were grouped by basidium, representing five unique basidia. Basidia groups where chosen for analysis if they contained three or more segregants with unique genotypes. Of the five basidia groups, two consisted of three segregants, two with four segregants, and one basidium exhibited eight unique genotypes.

#### Analysis of haplotypes with distorted allele frequencies

The allele frequency of haplotypes across segregants germinated from the same basidia was analyzed. Specifically, deviations from the expected 2:2 parental allele ratio where quantified. Regions were removed from consideration if only a single SNP supported the observation or if the size of the region was only one base in width. An ANOVA was used to examine average differences in size of haplotypes with distorted allele frequencies across the genome. A log-linear model was used to investigate the average number of haplotypes as a function of chromosome size (*α* = 0.05).

#### Analysis of allele bias

Across all the 39 segregants from the **a**-α bisexual crosses, a binomial model was used to identify chromosomal regions with bias towards one parental allele. This model assumed equal likelihood for inheriting either of the parental alleles (*p* = 0.50). SNP sites were collapsed across the 39 segregants based on recombination break points and common allele frequencies. This generated 944 sites to test in the binomial model. A false discovery rate approach fdr = 0.05) was used to correct for multiple comparisons. A similar procedure was used for testing for allele bias in the α-α unisexual cross.

### Data availability

Raw sequence reads generated form samples utilized in this study are available on NCBI’s sequence read archive under BioProject identification number PRJNA420966, with individual accession numbers SAMN08130857 – SAMN08130963. The generated variant call file from the aligned sequenced reads are publicly available on the GitHub repository https://github.com/magwenelab/crypto-recombination-paper.

## Results

*High density SNP data allows fine mapping of genome-wide crossovers*

Whole genome sequencing data was obtained for 55 segregants from α-α unisexual crosses between parental strains XL280αSS and 431α and 39 segregants from **a**-α bisexual crosses between the parental strains XL280**a** and 431α (Sun *et al.* 2014). Variants were called for each segregant (see Materials and Methods) and 86,767 biallelic, single nucleotide polymorphisms (SNPs) between the parental strains were used as genetic markers. Across the 19 Mb genome, comprised of fourteen chromosomes, the median distance between consecutive SNPs (inter-marker interval) was 87 bases with only 0.5% of the 86,753 inter-marker intervals larger than 2 kb (Supplementary Figure S1). SNP data was used to infer haplotypes and crossover events per segregant ((Figure 1). In total 3,301 crossovers were detected.

**Figure 1.**
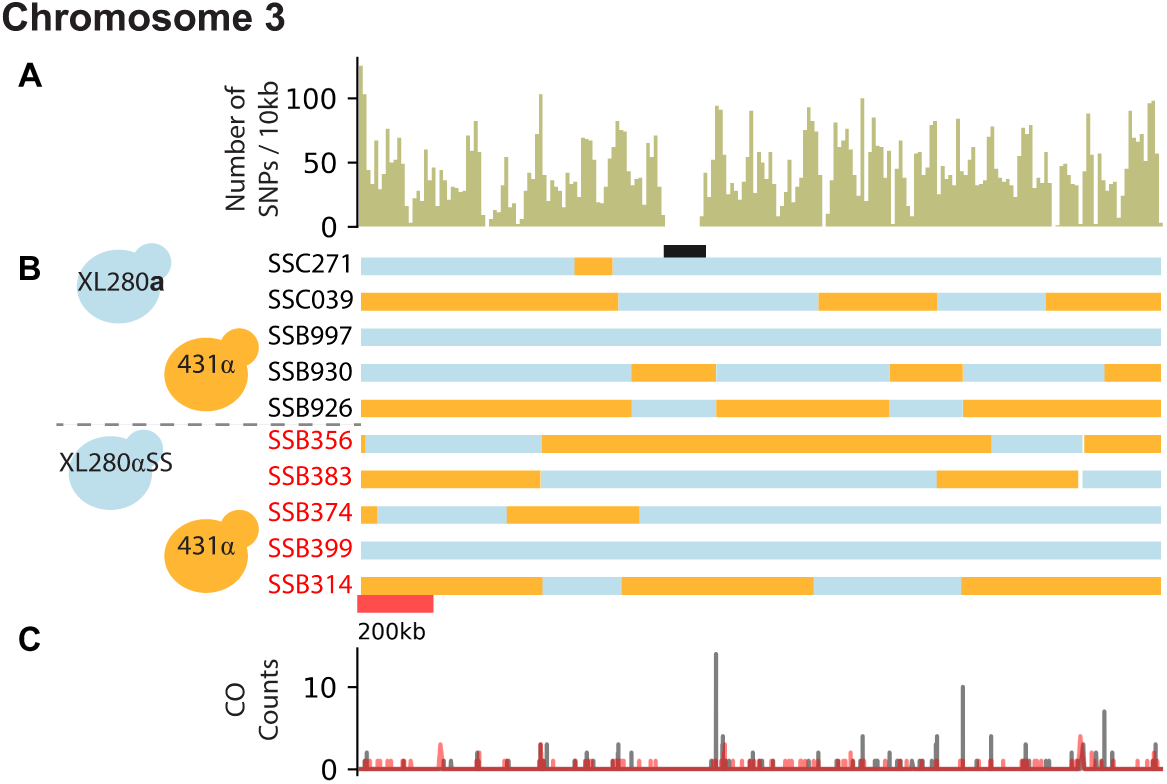
SNP density, haplotypes, and crossover counts of chromosome three. A) The SNP density for chromosome three (length ~2.1 Mb) across the progeny from the XL280**a** × 431α and XL280αSS × 431α crosses, calculated as the number of SNPs per 10 kb (total: 9,779 SNPs). B) Haplotypes, inferred from SNP data, are displayed as blue if inherited from XL280(αSS or a) or orange if inherited from 431α for 10 segregants from the α-α unisexual (red) and the **a**-α bisexual (black) crosses. The position of the centromere is displayed in black. C) Crossover (CO) counts along chromosome three for segregants from the α-α unisexual (red) and the **a**-α bisexual (black) crosses. Crossovers are detected by changes in genotype between two contiguous SNPs.

In each set of progeny from the α-α unisexual and **a**-α bisexual crosses, several segregants were identified as having at least one non-exchange chromosome. In 35 of 55 (64%) progeny from the α-α unisexual crosses and 19 of 39 (49%) progeny from the **a**-α bisexual crosses, at least one chromosome was non-recombinant based on filtered SNP data and inferred haplotypes. There is no difference in the distributions of number of non-exchange chromosomes per segregants across the two cross types (ks-test, p-value > 0.05). For these progeny, the median number of non-exchange chromosomes per segregant is between one and five. Smaller chromosomes are more likely to have zero crossovers. Of the 59 non-exchange chromosomes in the 35 progeny from the unisexual crosses, 32 (54%) have the parental XL280αSS genotype. However, in the 37 non-exchange chromosomes among the 19 progeny from the bisexual crosses, 29 (78%) have the XL280**a** parental copy.

### Genome-wide recombination rates differ between unisexual and bisexual reproduction in C. deneoformans

Genome wide recombination rates were estimated using Poisson regression, modeling the number of crossovers as a function of chromosome length with the mode of sexual reproduction as a covariate (see Materials and Methods). This model predicts an obligatory ~ 0.98 crossovers per chromosome for offspring from the unisexual crosses and ~ 1.30 crossovers per chromosome for offspring of the bisexual cross. There is a significant difference in the expected number of crossovers between segregants from α-α unisexual and **a**-α bisexual crosses (p-value < 10^−10^). The expected number of crossovers is predicted to increase by a ratio of ~ 1.768 per Mb increase in chromosome size ((Figure 2). Based on the sum of the per chromosome average and the total genome length, we estimate an approximate physical to genetic distance of 6.14 kb/cM for the α-α unisexual crosses and 4.67 kb/cM for the **a**-α bisexual crosses.

**Figure 2.**
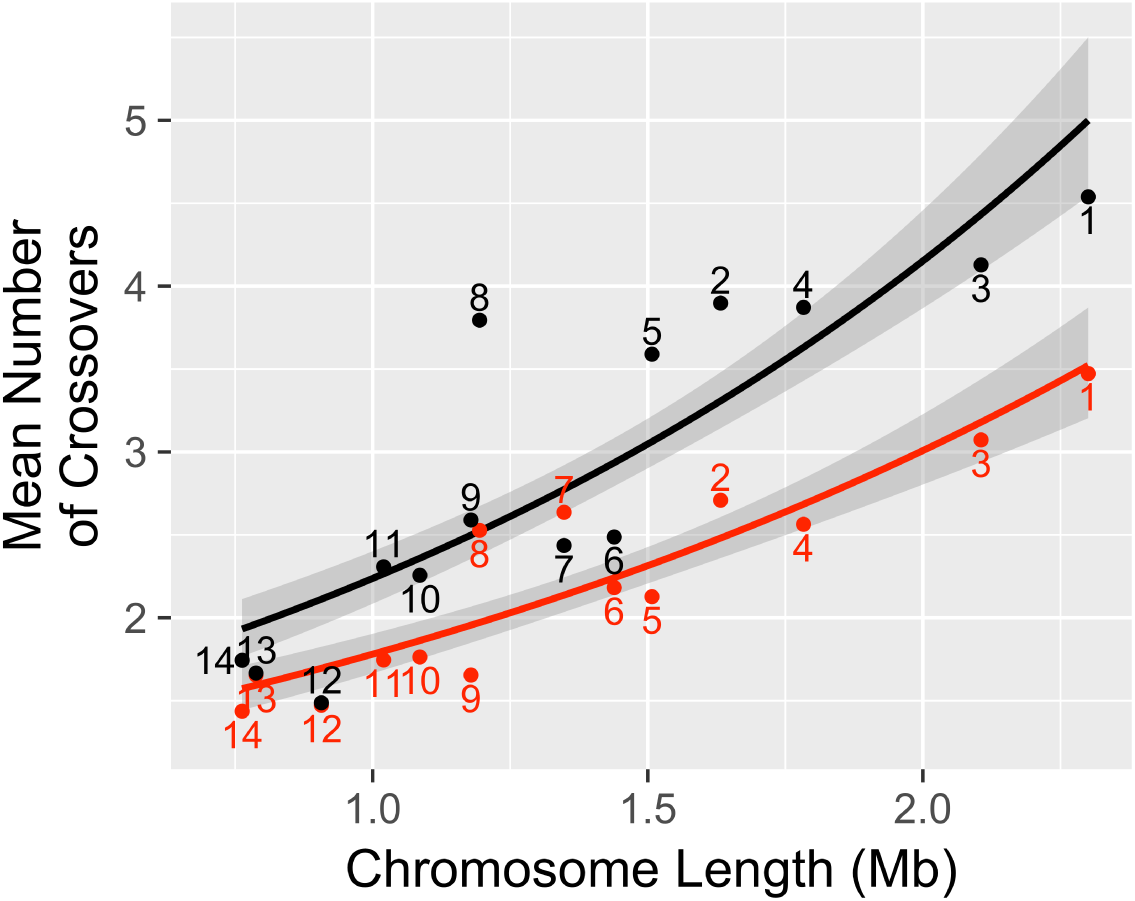
Unisexual vs bisexual crossovers as a function of chromosome length. The average number of crossovers for progeny from the α-α unisexual (red) and **a**-α bisexual crosses (black) are shown per chromosome. Solid lines indicate the estimated Poisson regressions for the two cross types separately, relating the number of crossovers to chromosome lengths. Shaded regions are 95% confidence intervals of the regression estimates. Numbers indicate chromosomes.

To explore this difference in greater detail, we compared recombination rates by chromosome for the two types of crosses. For chromosomes 1 – 5, 8, and 9 there are significant differences (Mann-Whitney U-test, q-values < 0.042) in the average number of detected crossovers between the progeny from the α-α unisexual and **a**-α bisexual crosses. No significant difference in the average number of crossovers between the two cross types was detected on chromosomes 6, 7, and 10-14 (Supplementary Figure S2).

### Analysis of crossover hot spots for segregants from α-α unisexual and **a**-α bisexual crosses in C. deneoformans

To identify crossover hot and cold spots along each chromosome, a binning approach was used. Bins of size 41.5 kb were tiled across each chromosome, and the number of crossovers detected within each bin was counted. The bin size of 41.5 kb was chosen based on simulations, so as to minimize the difference in the total number of hot and cold spots (see Materials and Methods). A Poisson model with this bin size and the expected genome-wide average crossover rate per segregant as estimated from the observed data (see Materials and Methods), was used in two tail tests to examine each bin for significantly high (hot) or low (cold) crossover rates. A false discovery rate procedure was used to establish genome-wide significance (*α* = 0.025, q-values < 0.014) (Benjamini and Yekutieli 2001). This analysis revealed 39 hot spots, bins with 20 or more detected crossovers, and 44 cold spots, bins with zero detected crossovers ((Figure 3). Along every chromosome, at least one crossover hot spot was identified and these regions are often found flanking or near centromeres.

**Figure 3.**
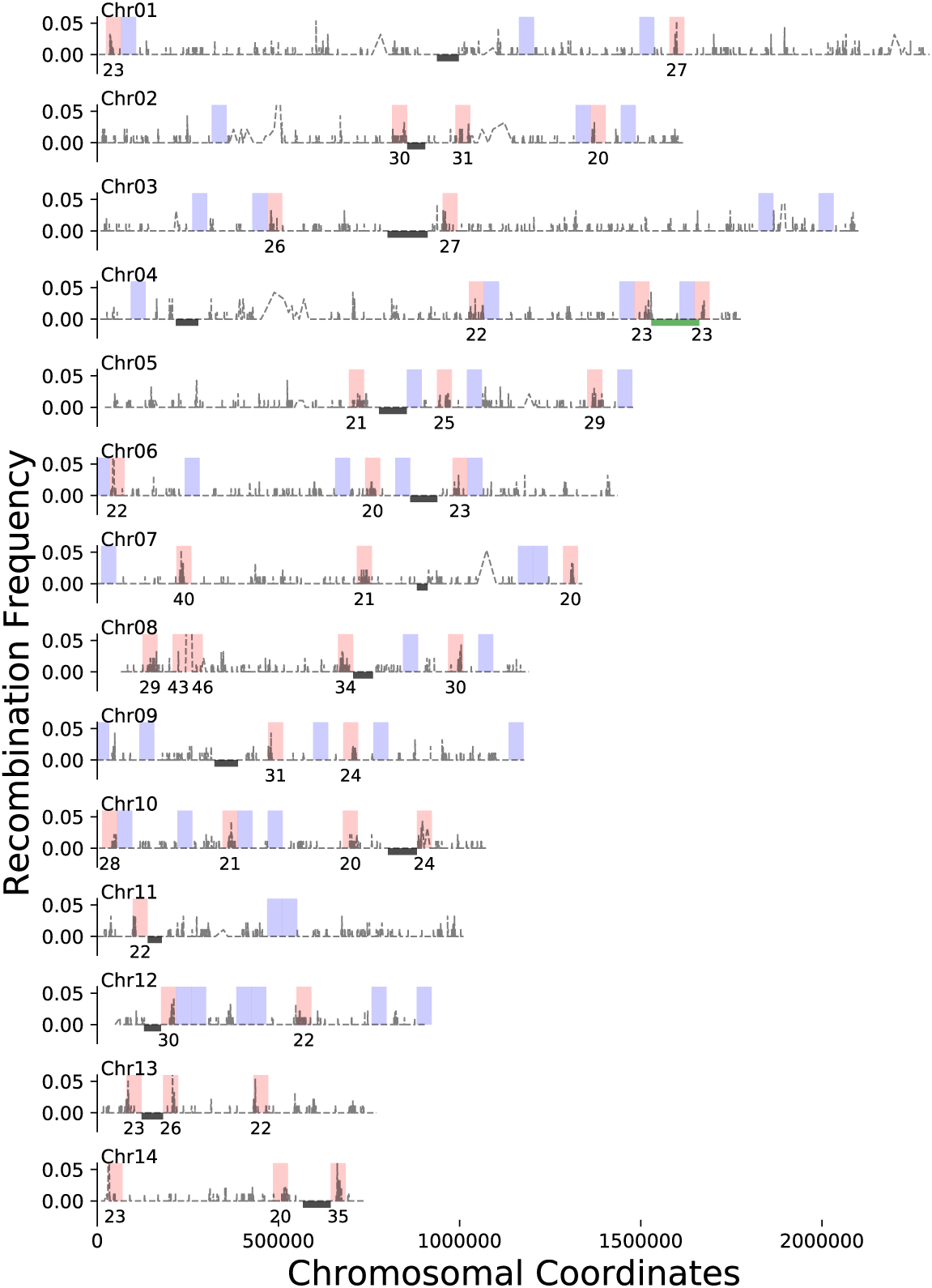
Genome-wide crossover hot and cold spots. In grey, the recombination frequencies (y-axis) for segregants from the α-α unisexual and **a**-α bisexual crosses along each of the fourteen chromosomes. Crossovers occur within an inter-marker interval and are detected as a change in genotype between consecutive SNPs. Bins, 41.5 kb wide, were used to segment each chromosome. For bins identified as crossover hot spots (red), the number of crossovers detected is labeled underneath. All crossover cold spots (blue) have zero detected crossovers. Locations of centromeres and the *MAT* locus are displayed as black bars and a green bar respectively. Note, the y-axis has been truncated in many instances to visualize crossovers along each chromosome.

Previous studies have demonstrated an association between recombination hot and cold spots and GC content (Sun *et al.* 2012; Sun and Heitman 2016). For 7,558 inter-marker interval sequences within the 39 hot spots, the mean GC content was ~0.49 (95% CI: [0.489, 0.494]) while the mean GC for 7,369 inter-marker interval sequences contained within the 44 cold spots was ~0.475 (95% CI: [0.473, 0.477]). The mean GC content of hot spots differs significantly from the cold spots (MannWhitney U-test, p-values < 10~^35^, Supplementary Figure S4). Both of these differ from the reported genome-wide average GC content (0.486) and the mean (~0.483, 95% CI: [0.482, 0.484]) of the other 68,051 inter-marker interval sequences not associated with hot or cold spots (Sun *et al.* 2012). Of the 7,558 inter-marker interval sequences within identified hot spots, 584 detect a genotype change (ie the approximate sites of double strand breaks) and of these inter-marker interval sequences, ~64.4% overlapped with intergenic regions when compared to the JEC21 annotation (Loftus *et al.* 2005).

From the set of 584 inter-marker interval sequences associated with hot spots and in which a crossover occurs, 100 random sequences were analyzed using MEME to identify sequence motifs associated with crossover hot spots. These sequences were compared to a control set of sequences selected in a similar fashion from other genomic regions. A poly(G) motif that is 29 bases long was identified in all of the 100 hot spot associated sequences (E-value < 10^−70^, Supplementary Figure S5).

### Allele bias and allele distortion seen in segregants generated via bisexual reproduction in C. deneoformans

Across the 39 segregants from the **a**-α bisexual cross, a binomial model was used to identify chromosomal regions with bias towards one parental allele, using a null model of equal likelihood of inheriting either of the parental alleles (*p* = 0.50). Five regions show evidence of biased allele inheritance towards the XL280**a** allele (q-value < 0.016). These regions are located on chromosomes one, two, four, six, and twelve with lengths of approximately 364, 260, 303, 41, and 60 kb respectively (Supplementary Figure S6). The allele frequencies across SNP sites in segregants from the α-α unisexual cross do not show evidence of bias towards either parental allele that reaches genome-wide significance.

Allele inheritance patterns within basidia were then examined for segregants from the **a**-α bisexual cross. Of the 39 progeny from the **a**-α bisexual crosses, 22 may be grouped by basidia of dissection. This grouping method generates five groups for analysis with three (*N* = 2), four (*N* = 2), and eight (*N* = 1) segregants, all with unique genotypes ((Figure 4). Using these segregants, representing five unique basidia, 197 regions were identified across the genome with allelic ratios deviating from the expected 2:2 (allelic distortion). The size of these regions with allelic distortion ranged from a minimum of six bases to a maximum of 1.4 Mb (Supplementary Figure S7, A). The average size of regions exhibiting allelic distortion does not differ across chromosomes (ANOVA, p-value = 0.092). The locations of allelic distortion regions are often similar across basidia ((Figure 4). Of the 197 allelic distortions, 83 were identified from basidia III, IV, and V with allele ratios consistent with possible gene conversions.

**Figure 4.**
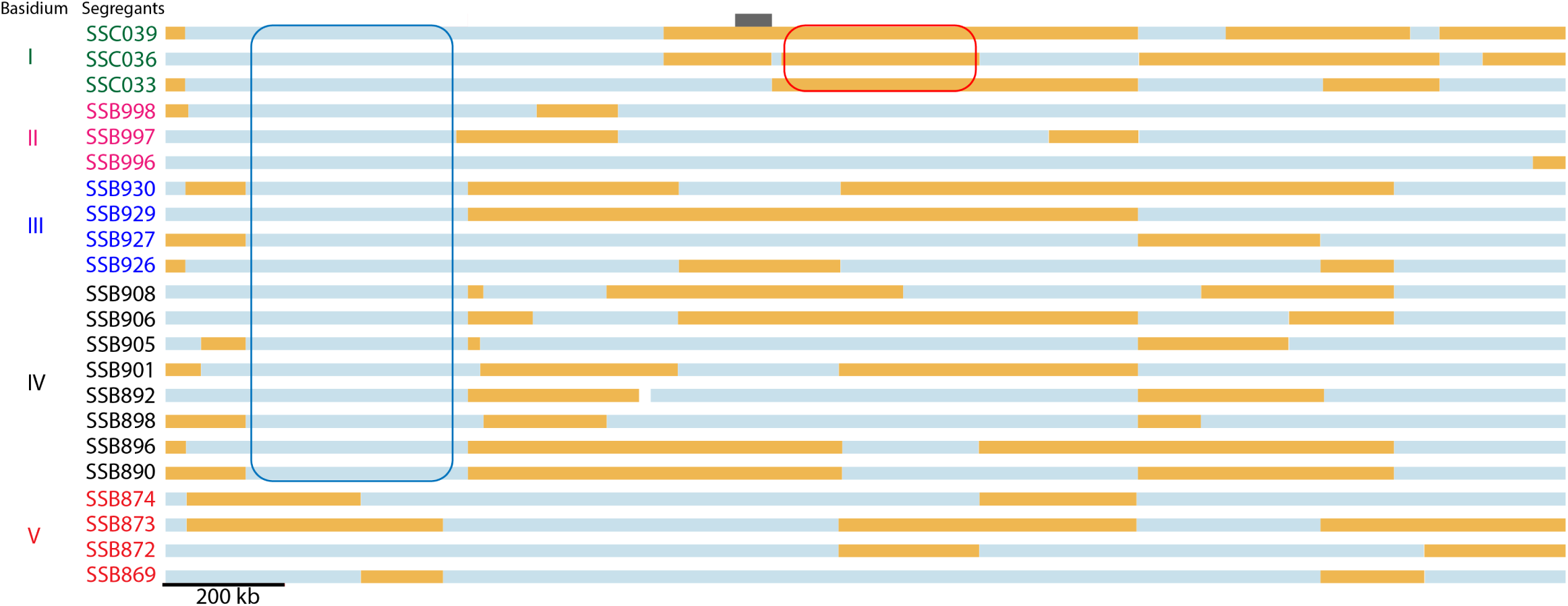
Allelic distortions along chromosome one in segregants from **a**-α bisexual crosses. Haplotypes (blue indicating inheritance from XL280**a**, orange from 431α) for 22 segregants from the **a**-α bisexual crosses, grouped by basidium of dissection. Circled in red is a region, in a single basidium, exhibiting allelic distortion in the direction of 431α. Circled in blue is a region that exhibits allelic distortion (towards XL280**a**) across multiple basidia. This second region overlaps with a region of allelic bias as determined from analysis of all progeny from the bisexual crosses. Other regions of allelic distortion are present in this figure. The position of the centromere is displayed as a black bar.

Across the 197 regions representing haplotypes with distorted parental allele frequencies, the specific allele inherited was examined. Along chromosome twelve, eleven haplotypes with distorted allele frequencies were identified, and ten of these retain the XL280**a** allele. Genome-wide, no evidence of consistent bias towards either parental genotype was observed (Supplementary Figure S7, B).

The average number of regions with distorted allele frequencies across the genome was established as a function of chromosome size for our 22 segregants representing 5 unique basidia from the **a**-α bisexual crosses (Supplementary Figure S8). A LogLinear model provides evidence supporting a significant association between chromosome size and the average number of haplotypes with distorted allele frequencies (p-value < 10^−5^).

### Unique patterns of allele segregation

Two groups of segregants from the **a**-α bisexual crosses representing two unique basidia showed interesting patterns of allele segregation. The first group of samples dissected from one basidium was comprised of eight spores and analysis of their recombinant haplotypes indicates all eight samples are genetically unique (for example see (Figure 4, basidium IV). This observation deviates from the expected four unique gametes expected to result from meiosis (Kwon-Chung 1980; Page and Hawley 2003; Idnurm *et al.* 2005). The second basidium showing interesting allele segregation was composed of four segregants. These four samples are all recombinant and were previously thought to be genetically unique as indicated by marker genotypes along chromosome four (Sun *et al.* 2014). However, our re-analysis indicates that two of the four segregants are nearly genetically identical; chromosome four is the only distinct chromosome differentiating the two samples, which are identical along the other thirteen chromosomes, including a partial duplication of chromosome ten.

## Discussion

*C. deneoformans* is capable of sexual reproduction between strains of the opposite and the same mating types. In this study we document higher rates of recombination in offspring generated from bisexual crosses. Progeny from the bisexual cross are predicted to have a basal rate of ~1.30 crossovers per chromosome versus ~0.98 crossovers per chromosome for progeny in the unisexual cross. For both sets of progeny, the number of crossovers is predicted to increase by a ratio of ~1.768 per Mb increase in chromosome size. Of the fourteen chromosomes in the *C. deneoformans* genome, seven show differences in the average number of crossovers per segregant when comparing samples from **a**-α bisexual and α-α unisexual crosses. Converting these crossover rates, we estimate an approximate physical to genetic distance of 6.14 and 4.67 kb/cM for the α-α unisexual and **a**-α bisexual crosses, respectively. These estimates are nearly three times lower than the estimated crossover rate of *Saccharomyces cerevisiae* (~2 kb/cM, Cherry *et al.* (1997); Barton *et al.* (2008)) and far higher than the crossover rates estimated for *Drosophila melanogaster* (~100 kb/cM, *Comeron et al.* (2012)), *Arabidopsis thaliana* (~278 kb/cM, Salomé *et al.* (2012)), and *Homo sapiens* (~840 kb/cM, Kong *et al.* (2002)).

Our results differ from previous estimates because they are based on information from the entire *C. deneoformans* genome and utilize more than 200 fold higher density of markers than have been employed in any previous study of recombination in *C. deneoformans* (Forche *et al.* 2000; Marra *et al.* 2004; Sun *et al.* 2014). For example, relative to the earlier study of Sun *et al.* (2014), which utilized the same set of offspring, we detected differences in the average number of crossovers along chromosome four between progeny from α-α unisexual and **a**-α bisexual crosses. We reasoned that this difference was due to increases in the detected number of crossovers resulting from increased marker density. To confirm this, SNPs were selected to best approximate marker locations from Sun *et al.* (2014) such that the maximum difference in location between these SNPs and the previous marker location was one kilobase. Using these data to reconstruct haplotypes and calculate crossover events recapitulated the previous findings. Thus differences in observed recombination events along chromosome four, relative to a prior report that analyzed the same segregants, are due to increased marker density which facilitates the detection of genotype changes previously masked by double crossover events (Supplementary Figure S3).

The regression model used to relate chromosome length and the number of crossovers predicts nearly one obligate crossover *on average* per chromosome for both sets of progeny from the α-α unisexual and **a**-α bisexual crosses (see Results). A significant number of segregants had chromosomes that had zero detected crossovers (non-exchange chromosomes), but analysis of segregants from basidia groups suggests that the standard model of crossover assurance holds (i.e. there is at least one crossover per homologous chromosome pair per meiosis; Ault and Nicklas (1989)). The non-exchange chromosomes we observed may thus be due to Holiday junctions resolving into noncrossover events during chromosome disjunction or may reflect chromatids that weren’t involved in crossovers during meiosis.

The analysis of crossover hot and cold spots identified at least one crossover hot spot along each of the fourteen chromosomes, and cold spots on every chromosome except 13 and 14. Analyses based on a subset of the hot spot inter-marker interval sequences in which crossovers were detected, identified a poly(G) motif significantly enriched within these sequences. Furthermore, inter-marker interval sequences within crossover hot spots have on average higher GC content, as documented in other studies of *C. deneoformans* as well as other fungi (Gerton *et al.* 2000; Petes 2001; Mancera *et al.* 2008; Marsolier-Kergoat and Yeramian 2009; Sun *et al.* 2012; Sun and Heitman 2016). Of the crossover hot spots, two were identified that flank the *MAT* locus, recapitulating the findings of several other studies (Marra *et al.* 2004; Hsueh *et al.* 2006; Sun *et al.* 2012; Sun and Heitman 2016). While recombination hot spots flank the *MAT* locus, the *MAT* locus itself contains a crossover cold spot, consistent with previous findings (Sun *et al.* 2014). Parallel to the pattern observed at the *MAT* locus, we noted a tendency for centromeric regions and crossover cold spots tend to be surrounded by flanking crossover hot spots. Some caution is required in interpreting the total number of hot and cold spots, and their precise locations. Due to the SNP and haplotype filtering criteria we employed, some genomic regions such as centromeres and telomeres are excluded from analysis. Thus we are unable to access recombination or gene conversion events that could have taken place within centromeric regions, as suggested in previous studies of *Cryptococcus* (Janbon *et al.* 2014; Sun *et al.* 2017) and other fungal species such as *Candida albicans* (Thakur and Sanyal 2013). The precise location of inferred hot and cold spots is also a function of the choice of bin widths and starting coordinates.

In addition to providing genome-wide information on crossover hot and cold spots, our analysis identified numerous regions that have allele ratios that deviate from the expected 2:2 parental ratio in progeny from the **a**-α bisexual crosses, consistent with the findings of Sun *et al.* (2014) for chromosome four. Some of the regions with deviant allele frequencies have 3:1 allele ratios which would be consistent with gene conversion, but most of the regions of allelic distortion are quite large, nearing 100 kb. Thus it is unlikely that gene conversions alone explain the observed loss of heterozygosity genome-wide, as conversion tracks from gene conversions are thought to be small, on the order of only a few kilobases as observed in *S. cerevisiae* (Mancera *et al.* 2008). Alternate models that could explain the observed allelic distortions include mitotic recombination that takes place after nuclear fusion but prior to meiosis, or chromosomal mis-segregation that takes place during cell fusion prior to meiosis and formation of a basidium (leading to loss of a parental genotype). Chromosomal breakage prior to meiosis and then repair using the homologous chromosome could also lead to a loss of one of the parental alleles (Sun *et al.* 2014).

Of the segregants from the **a**-α bisexual crosses, two groups are worth discussing in detail. The first group is comprised of four segregants from a single basidium. All four segregants were previously described as unique based on marker genotypes along chromosome four (Sun *et al.* 2014). However, genome-wide analysis revealed that two of the segregants are genetically identical except for chromosome four and are aneuploid for chromosome ten. For this set of segregants the patterns of allele segregation could be explained by chromosomal non-disjunction. During the formation of the basidium and during meiosis, chromosomal non-disjunction could have produced three nuclei, two with the correct ploidy of both chromosome four and ten and one nucleus with two unique, recombinant copies of chromosome four. Such patterns have been observed in hybrid crosses between *C. neoformans* and *C. deneoformans* (Vogan *et al.* 2013). During mitosis and basidiospore packaging, this aneuploid nucleus may have produced several copies of itself with varying arrangements of the genome, thus generating haplotypes genetically identical except for chromosome four as seen in two of these segregants. The aneuploidly of the tenth chromosome in these segregants can be explained by the known aneuploidy of this chromosome in the parental strain XL280**a** (Sun *et al.* 2012). Another basidium from the **a**-α bisexual crosses that exhibited interesting patterns of allele segregation was a collection of eight segregants. Analysis of the haplotypes of these eight segregants indicates all are genetically unique. In this instance, fusion between sister haploid nuclei could have taken place post meiosis within the basidium, providing opportunity for mitotic recombination to occur and, through subsequent rounds of mitosis, produce more than four unique gametes (Vogan *et al.* 2013). Due to the nature of *C. deneoformans* and the methods of dissection, it is almost impossible to determine if crossover events occur during meiosis or mitosis.

Our analyses provide evidence of different rates of recombination in unisexual and bisexual crosses, however the mechanisms that drive such differences are as yet unknown. Could these differences be mating type specific? While the gene contents between the *MAT***a** and *MAT*α alleles are similar, mating type specific regulators such as the heterodimeric transcription factor *SXI1*α/*SXI2***a**, are known to regulate a variety of processes involved in diploid sexual development (Hull *et al.* 2005; Mead *et al.* 2015). Here we postulate that the presence or absence of mating type specific factors may change the regulation of genes critical for recombination, such as *DMC1* and *SPO11* (Lin *et al.* 2005), leading to higher or lower crossover rates during sexual reproduction.

In this report we have focused on a single species, and the extent to which the patterns and rates of recombination we document here for *C. deneoformans* hold across all *Cryptococcus* species and lineages is as yet unknown. Like *C. deneoformans*, in the VNI and VNII lineages of *C. neoformans* most isolates are of the *MAT*α mating type (Kwon-Chung and Bennett 1978). Only in populations of the VNBI and VNBII lineages are *MAT***a** strains found with significant frequency (Litvintseva *et al.* 2003; Desjardins *et al.* 2017). This has led to the hypothesis that sexual reproduction in many *C. neoformans* lineages may be primarily unisexual (Fu *et al.* 2015). The differences in rates of recombination we document here between **a**-α bisexual and α-α unisexual matings may contribute to differences in population recombination rates, even if **a**-α bisexual and α-α unisexual matings occur at similar frequencies. Consistent with this idea, the analysis of Desjardins *et al.* (2017) indicates that linkage disequilibrum decays at a relatively similar rate in both VNB lineages (bisexual) and the VNI lineage (unisexual). However, the primarily unisexual VNI lineage shows an overall higher rate of linkage disequilibrium. New high resolution genomic data, both from crosses and from population studies (Desjardins *et al.* 2017; Rhodes *et al.* 2017), will help to clarify the relative contributions that sex, mitotic recombination (Vogan *et al.* 2013), hypermutation (Billmyre *et al.* 2017), and other mechanisms for generating genomic variation contribute to the origins and maintenance of genetic diversity within this clade of fungal pathogens.

## Acknowledgments

This research was funded by the Research Training Grant from the University Program in Genetics and Genomics, Duke University, the Office of Biomedical Graduate Diversity’s IMSD program, BioCoRE, NIH grants R56AI123502, R01AI133654, and R37AI39115-20. We would also like to thank Dr Debra Murray and Dr Selcan Aydin for comments and feedback on the manuscript.

## Author Contributions

Conceived and designed the experiments: SS JH. Performed the experiments: SS. Analyzed the data: CR PMM. Contributed reagents and materials: SS RBB JH. Wrote the paper: CR PMM. Edited the paper: CR SS RBB JH PMM.

## Supplementary Figures

**Figure S1.**
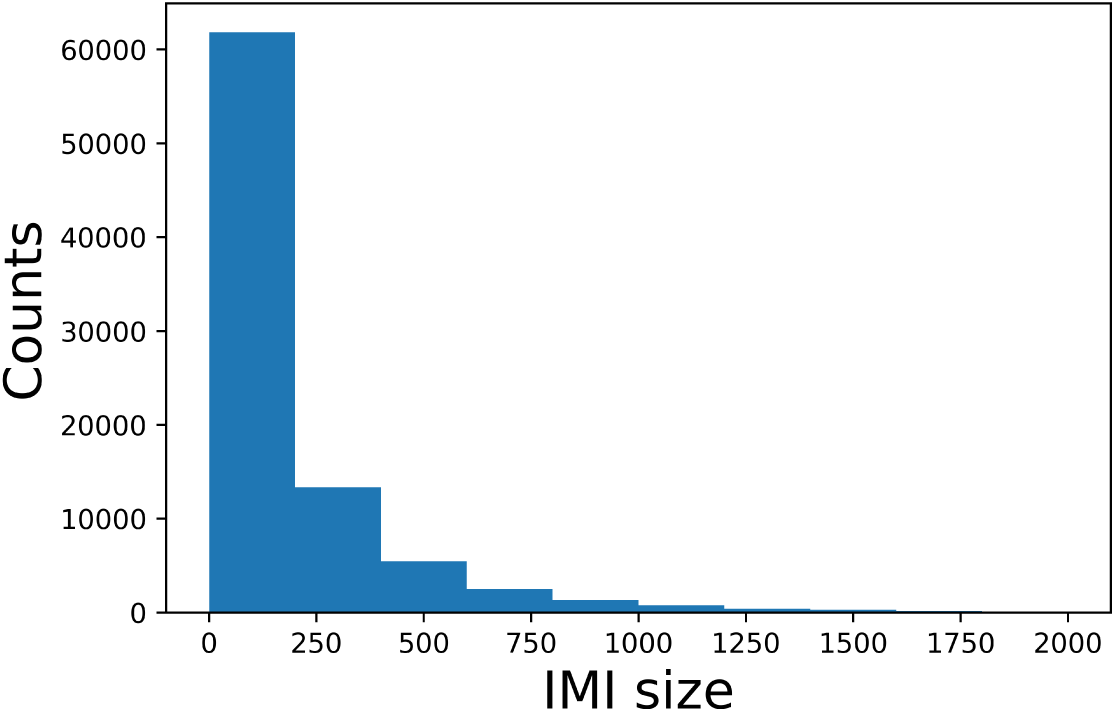
Distribution of inter-marker interval size across progeny from the from the XL280**a** × 431α and XL280αSS × 431α crosses. The total number of inter-marker intervals is 86,753. There are 86,278 inter-marker intervals with size < 2 kb. Only 0.548% of the inter-marker intervals have a size greater than 2 kb (data not shown). The median inter-marker interval size is 87 bases.

**Figure S2.**
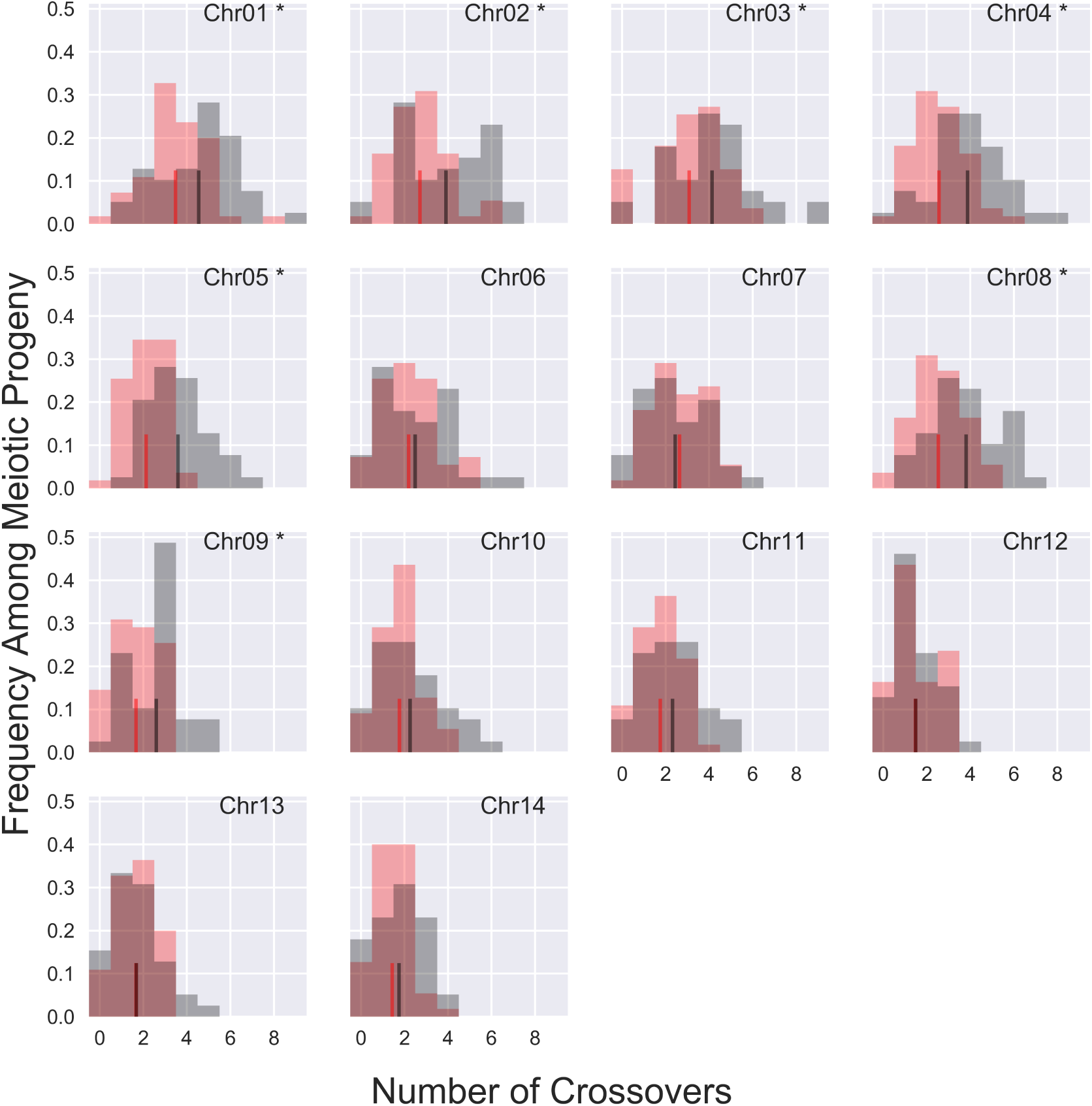
Distributions of crossovers per chromosome. The means of each distribution are displayed as red and black vertical lines for the segregants from the unisexual and bisexual crosses, respectively.“*” indicates chromosomes that show significant difference in the mean number of crossovers per segregant between progeny from the α-α unisexual unisexual (red) and **a**-α bisexual bisexual (black) crosses.

**Figure S3.**
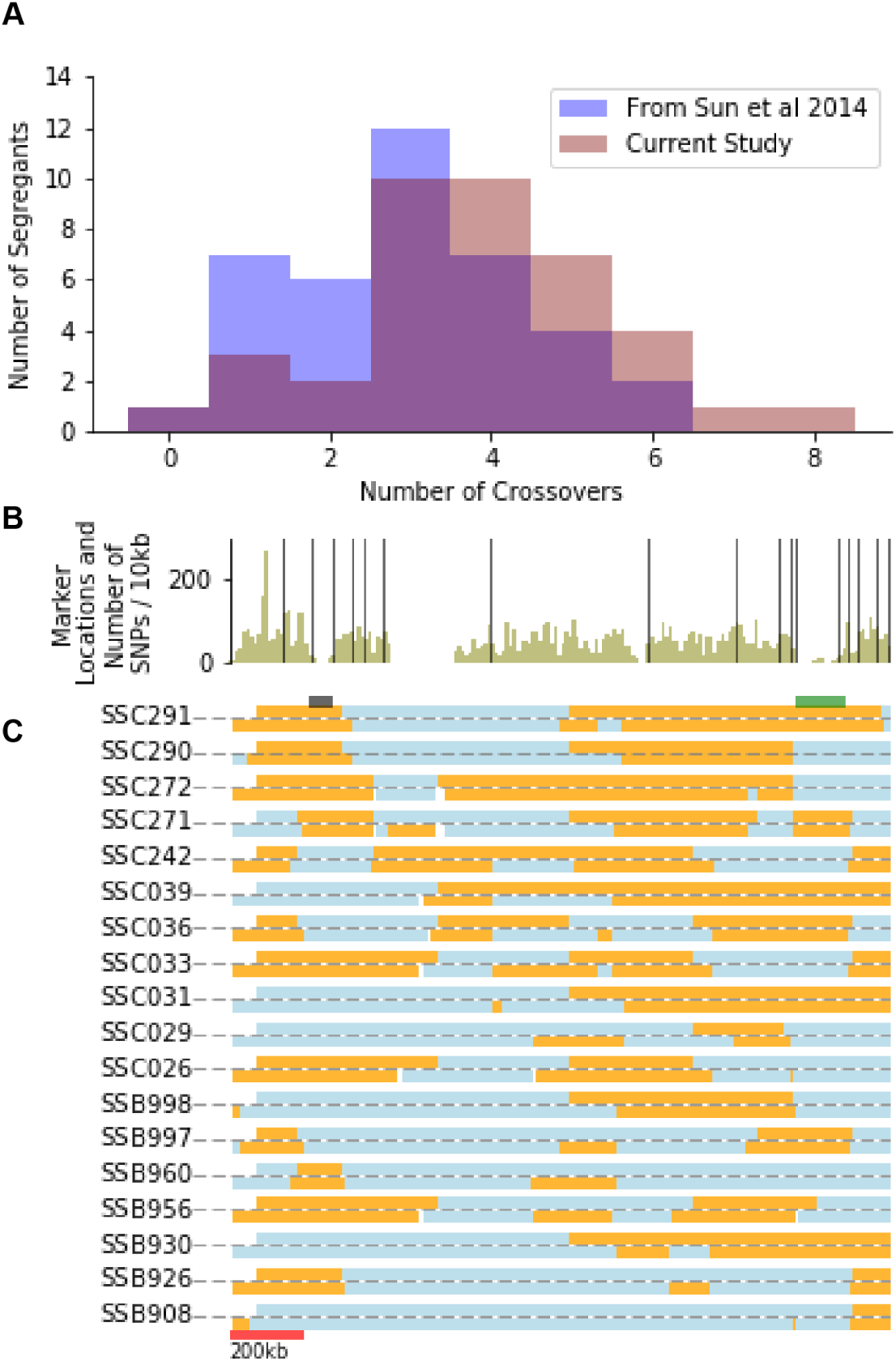
Changes in detected crossover along chromosome four are due to increased marker density. A) Distributions of crossovers along chromosome four for segregants from the **a**-α bisexual bisexual crosses. Recapitulated crossover counts from Sun *et al.* (2014) are shown in dark blue and current counts in red. B) Marker locations and SNP density across chromosome four. Locations of SNPs used to recapitulate results from Sun *et al.* (2014) are shown as solid black, vertical lines. The SNP density every 10 kb is shown in green. C) Inferred haplotypes from SNP data for segregants from the **a**-α bisexual crosses with detected differences be-tween the current study and Sun *et al.* (2014) For each segregant, the haplotype inferred from SNPs near marker locations used in Sun *et al.* (2014) (above grey line) and haplotypes from SNP data generated in this study (below grey line) are shown. Blue regions represent genetic material inherited from the XL280**a** parental strain. The approximate locations of the centromere and *MAT* locus are shown as black and green bars, respectively.

**Figure S4.**
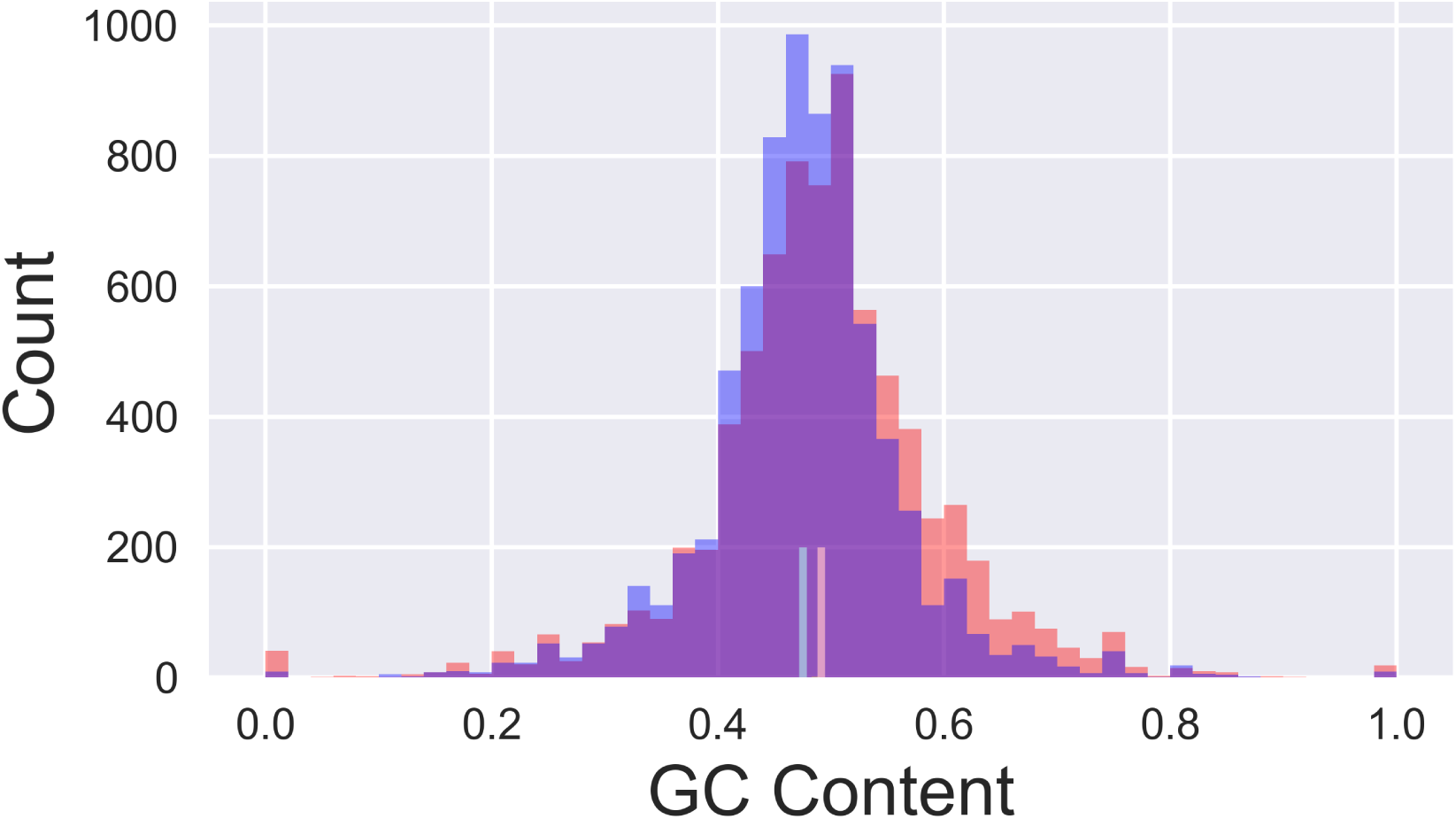
Distributions of GC content for sequences associated with recombination hot (red) and cold (blue) spots. Vertical lines show mean GC content for sequences associated with recombination hot (red) and cold (blue) spots.

**Figure S5.**
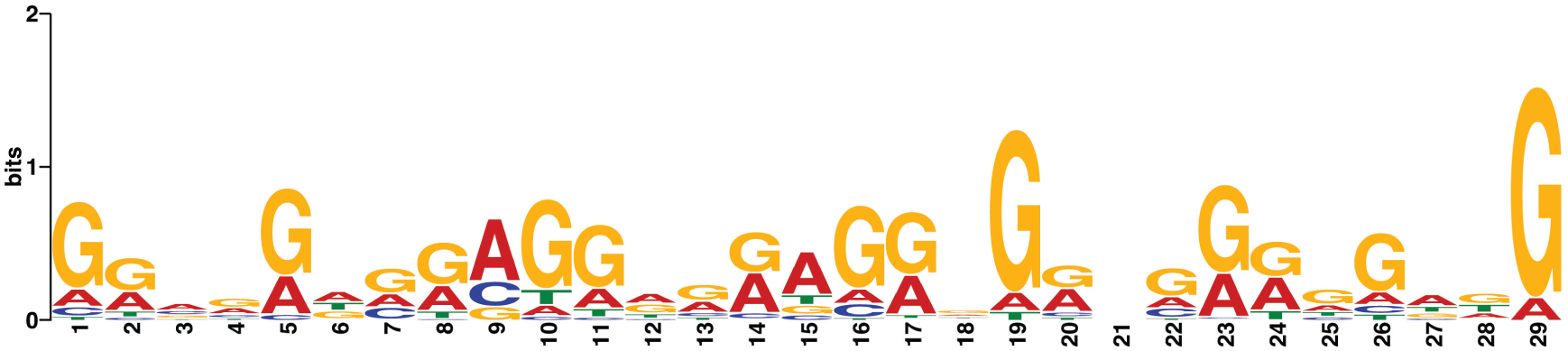
Poly(G) motif associated with crossover hot spots. This motif was found in all of the randomly chosen 100 inter-marker interval sequences associated with crossover hot spots submitted to MEME.

**Figure S6.**
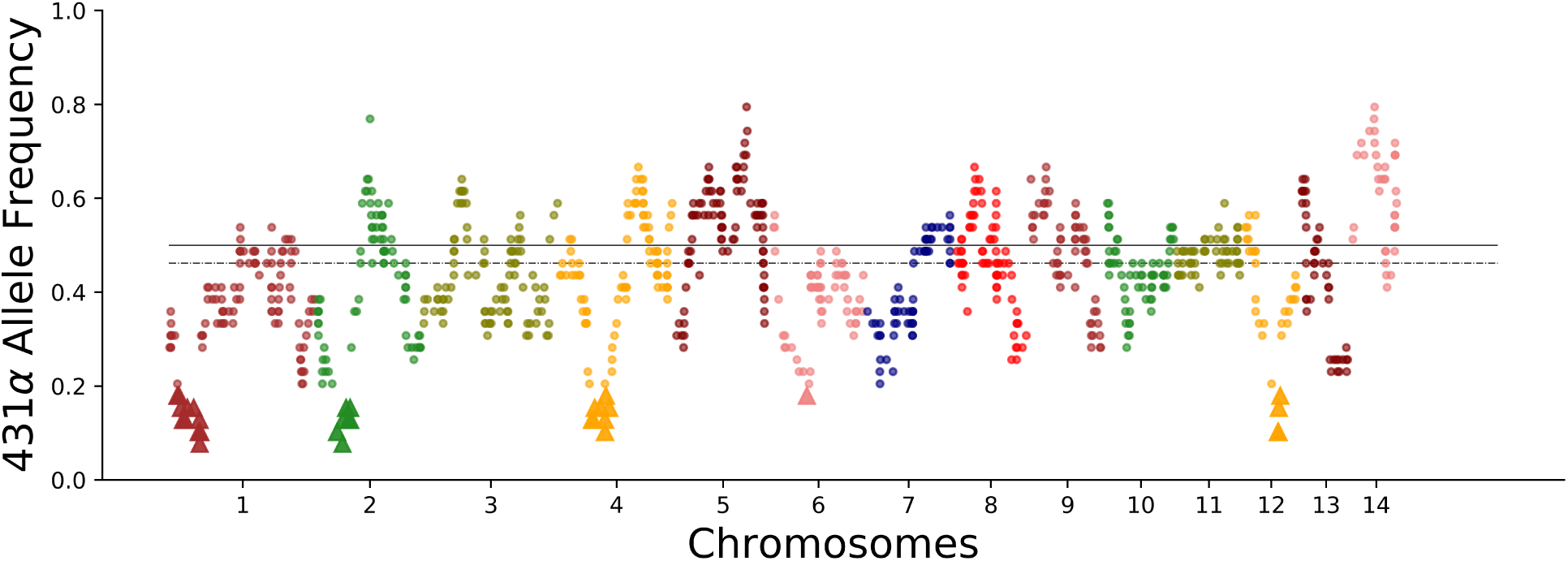
Allele bias in segregants from bisexual crosses. The genome-wide frequencies of the 431α parental allele in the 39 progeny from the **a**-α bisexual crosses. Triangles denote five regions along chromosomes one, two, four, six, and twelve with lengths of ~ 364, 260, 303,41, and 60 kb, respectively, biased towards the XL280**a** parental allele. Solid and dashed lines indicate an allele frequency of 0.5 and the median, genome-wide allele frequency of 0.46, respectively.

**Figure S7.**
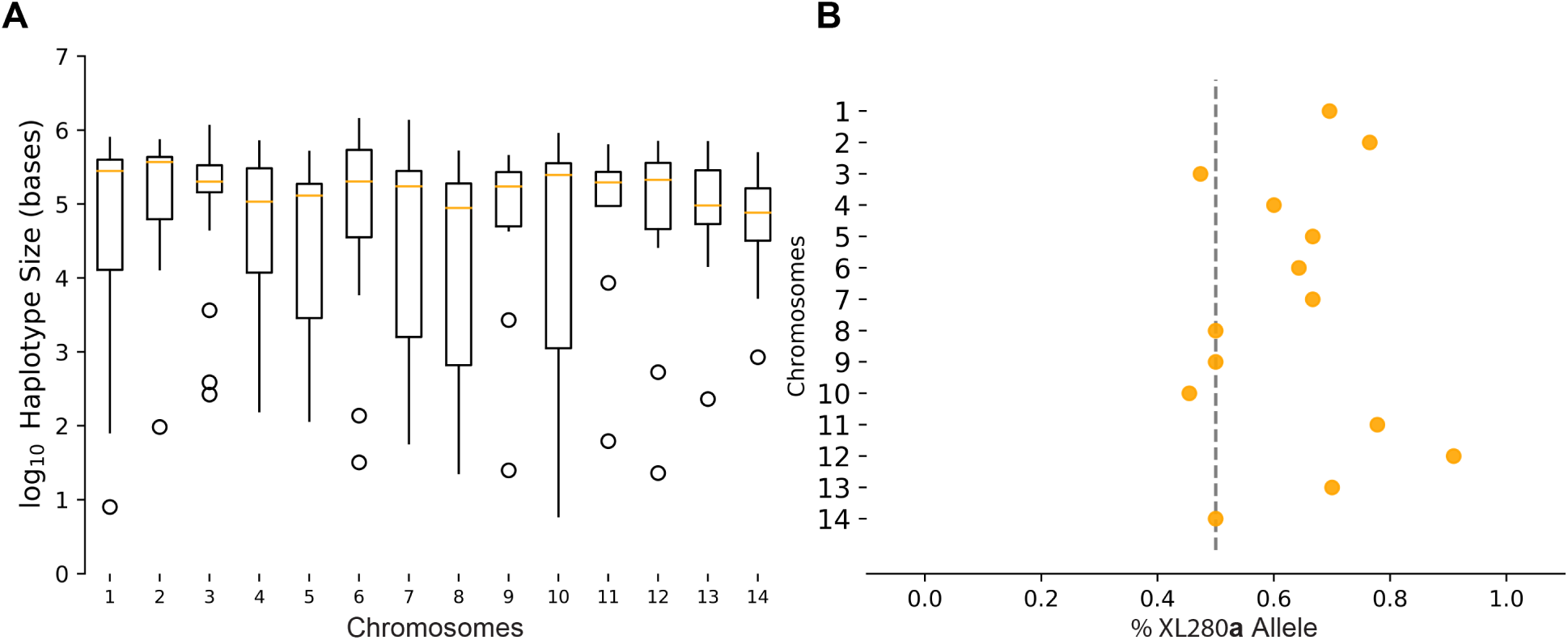
Size of haplotypes deviating from the expected 2:2 parental allele ratio. A) the log_10_ of haplotype size with distorted allele frequencies per chromosome. B) the percentage of haplotypes with distorted allele frequencies with the XL280**a** parental allele per chromosome.

**Figure S8.**
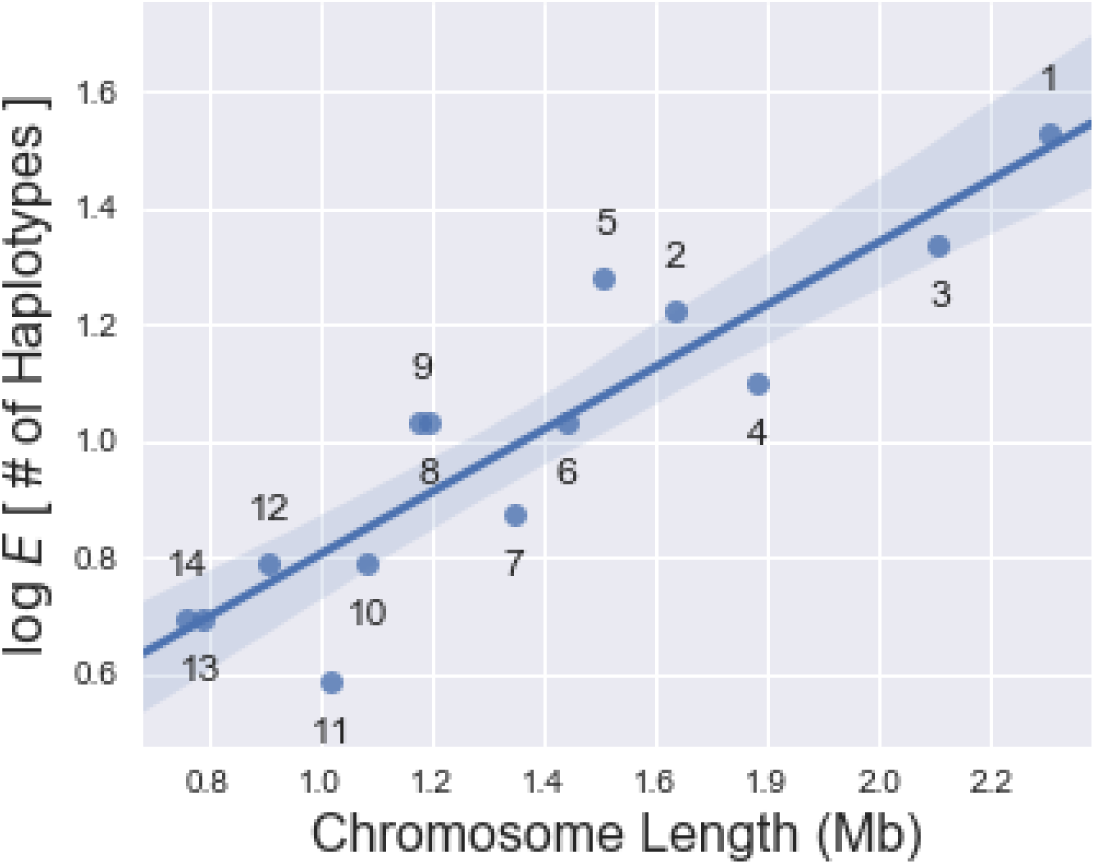
Genome-wide analysis of distorted haplotypes. The log of the average number of haplotypes within a basidium with allele frequencies deviating from the expected 2:2 parental ratio as a function of chromosome length is shown. The blue line represents a log-linear model, shaded regions represent the 95% confidence interval for regression estimates. Numbers dictate chromosomes.

